# *Pseudomonas aeruginosa* uses c-di-GMP phosphodiesterases RmcA and MorA to regulate biofilm maintenance

**DOI:** 10.1101/2020.06.26.174680

**Authors:** S.L. Katharios, G.B. Whitfield, P.L. Howell, G.A. O’Toole

## Abstract

While the early stages of biofilm formation have been well characterized, less is known about the requirements for *Pseudomonas aeruginosa* to maintain a mature biofilm. We utilized a *P. aeruginosa*-phage interaction to find that *rmcA* and *morA*, two genes which encode for c-di-GMP-degrading phosphodiesterase (PDEs) enzymes, are important for the regulation of biofilm maintenance. Deletion of these genes initially results in an elevated biofilm phenotype characterized by increased production of c-di-GMP, Pel polysaccharide and biofilm biomass. In contrast to the wild-type strain, these mutants were unable to maintain the biofilm when exposed to carbon-limited conditions. The susceptibility to nutrient limitation, and subsequent loss of biofilm viability of these mutants, was phenotypically reproduced with a stringent response mutant (Δ*relA* Δ*spoT*), indicating that the Δ*rmcA* and Δ*morA* mutants may be unable to appropriately respond to nutrient limitation. Genetic and biochemical data indicate that RmcA and MorA physically interact with the Pel biosynthesis machinery, supporting a model whereby unregulated Pel biosynthesis contributes to the death of the Δ*rmcA* and Δ*morA* mutant strains in an established biofilm when nutrient-limited. These findings provide evidence that c-di-GMP-mediated regulation is required for mature biofilms of *P. aeruginosa* to effectively respond to changing availability of nutrients. Furthermore, the PDEs involved in biofilm maintenance are distinct from those required for establishing a biofilm, thus indicating that a wide variety of c-di-GMP metabolizing enzymes in organisms like *P. aeruginosa* likely allows for discrete control over the formation, maintenance or dispersion of biofilms.

**Importance:** Recent advances in our understanding of c-di-GMP signaling have provided key insights into the regulation of biofilms. Despite an improved understanding of how they initially form, the processes that facilitate the long-term maintenance of these multicellular communities remain opaque. We found that *P. aeruginosa* requires two phosphodiesterases, RmcA and MorA, to maintain a mature biofilm and that *P. aeruginosa* biofilms lacking these PDEs succumb to nutrient limitation and die. The biofilm maintenance deficiency observed in Δ*rmcA* and Δ*morA* mutants was also found in the stringent response defective Δ*relA* Δ*spoT* strain, suggesting that a regulatory intersection between c-di-GMP signaling, EPS biosynthesis and the nutrient limitation response is important for persistent surface growth. We uncover components of an important regulatory system needed for *P. aeruginosa* to persist in nutrient-poor conditions, and provide some of the first evidence that maintaining a mature biofilm is an active process.

## Introduction

*Pseudomonas aeruginosa* is a Gram-negative opportunistic pathogen that is found both environmentally and within clinical settings. Able to transition from planktonic lifestyles to a biofilm mode of growth, *P. aeruginosa* biofilms develop via a number of discrete steps generally defined as initial attachment, irreversible attachment, microcolony formation, maturation and dispersal (1). Flagella mediate the initial polar attachment of the cell to the surface, while pili facilitate irreversible attachment and commitment to surface growth (2, 3). Once on the surface, increased production of extracellular polysaccharide (EPS) facilitates increased surface adhesion, intercellular cohesion and provides both protection and structural integrity for mature biofilms to form (3–6).

An important element in the transition of *P. aeruginosa* from a planktonic to biofilm mode of growth is bis-(3′,5′)-cyclic dimeric guanosine monophosphate (c-di-GMP), a second messenger that coordinates the regulatory control of virulence and behaviors needed for surface growth such as motility and the production of EPS (7–10). The concentration of intracellular c-di-GMP is controlled enzymatically by c-di-GMP synthesizing diguanylate cyclases (DGCs) which contain GGDEF domains, and c-di-GMP degrading phosphodiesterases (PDEs) that harbor EAL or HD-GYP domains (8, 11). The genome of *P. aeruginosa* PA14 encodes ∼40 different DGCs and PDEs involved in the regulation of c-di-GMP (12). Despite the published record of the contributions of these different enzymes to biofilm formation and dispersion (13, 14), our understanding of how they are coordinated and integrated temporally to affect bacterial behavior remains incomplete.

Select DGCs and PDEs have been shown to be important during different phases of biofilm formation. For example, the DGC SadC and the PDE BifA both contribute to irreversible attachment and early biofilm formation (15, 16). During subsequent biofilm growth, the DGC WspR mediates biofilm maturation by increasing the production of EPS (17). Regulation of c-di-GMP also facilitates the return to the planktonic lifestyle, as evidenced by the PDEs NbdA and DipA, which are required for dispersion in response to changes in nutrients and nitric oxide levels, respectively (18, 19).

While available literature provides substantial insights into how biofilms form and disperse, our understanding of biofilm maintenance - the process by which existing biofilms regulate themselves to persist on a surface is rudimentary. Indeed, it is not even clear if maintenance of the biofilm is an active process. To date, information regarding biofilm maintenance is largely informed by proteomic analysis of biofilms at specific stages of development (20), an analysis primarily conducted using biofilms grown under nutrient-rich steady-state conditions (20), leaving open the questions of if and how the regulation of biofilms occurs during starvation. Here, we provide evidence that the PDEs RmcA and MorA are needed for the maintenance of *P. aeruginosa* biofilms - the loss of either of these PDEs results in increased EPS production and biomass in nutrient-sufficient conditions, but increased cell death and compromised biofilms during starvation.

## Results

### CRISPR-activated genetic background reveals role of c-di-GMP signaling in biofilm maintenance

Previously, we reported that the chromosomal integration of a 42-nucleotide sequence of DNA from the bacteriophage DMS3 into the genome of *P. aeruginosa* PA14 resulted in a CRISPR-activated genetic background. The CRISPR-activated strain that carries this 42-nucleotide sequence in the attachment (*att*) site, called *att*::DMS3_42_, was reported to be biofilm-negative due to increased cell death after 24 h, a typical time point to assess biofilm formation in a standard 96 well, static biofilm assay (21). Additionally, while the *att*::DMS3_42_ strain was able to form a biofilm at early time points (∼6 h) it was largely biofilm-negative by 12 h, suggesting that biofilm formed but could not be maintained. These data indicated to us that *att*::DMS3_42_ could serve as an effective tool to probe for functions required for biofilm maintenance, a poorly characterized aspect of the biofilm life cycle. Our goal then was to exploit the robust phenotype offered by the CRISPR-activated strain to study biofilm biology.

Due to the important role of c-di-GMP in facilitating surface growth, we hypothesized that reduced levels of this second messenger contributed to the inability of *att*::DMS3_42_ to maintain a biofilm at later time points. To test this hypothesis, we employed a vector expressing the GcbC-R363E mutant protein, which codes for a DGC found in *Pseudomonas fluorescens* that has a mutation in the regulatory I-site that renders it constitutively active, thereby enhancing c-di-GMP synthesis (22). We heterologously expressed this construct in the *P. aeruginosa* PA14 WT and *att*::DMS3_42_ strain backgrounds to assess whether increased c-di-GMP synthesis could rescue the biofilm deficiency observed at 12 h for the latter strain. Introduction of the *gcbC*-R363E expression plasmid into the WT strain resulted in elevated biofilm biomass over the empty vector control, consistent with the increased production of c-di-GMP (**Fig. 1A**). Consistent with our hypothesis, the expression of this constitutively-active DGC in the strain carrying the att::DMS3_42_ construct greatly reduced the biofilm defect (i.e., enhanced biofilm formation) at 12 h compared to the att::DMS3_42_ strain carrying the empty vector (EV) control (**Fig. 1A**). These data suggest that the inability of the strain carrying the *att*::DMS3_42_ insertion to maintain the biofilm may be the result of decreased levels of c-di-GMP.

**Figure 1.**
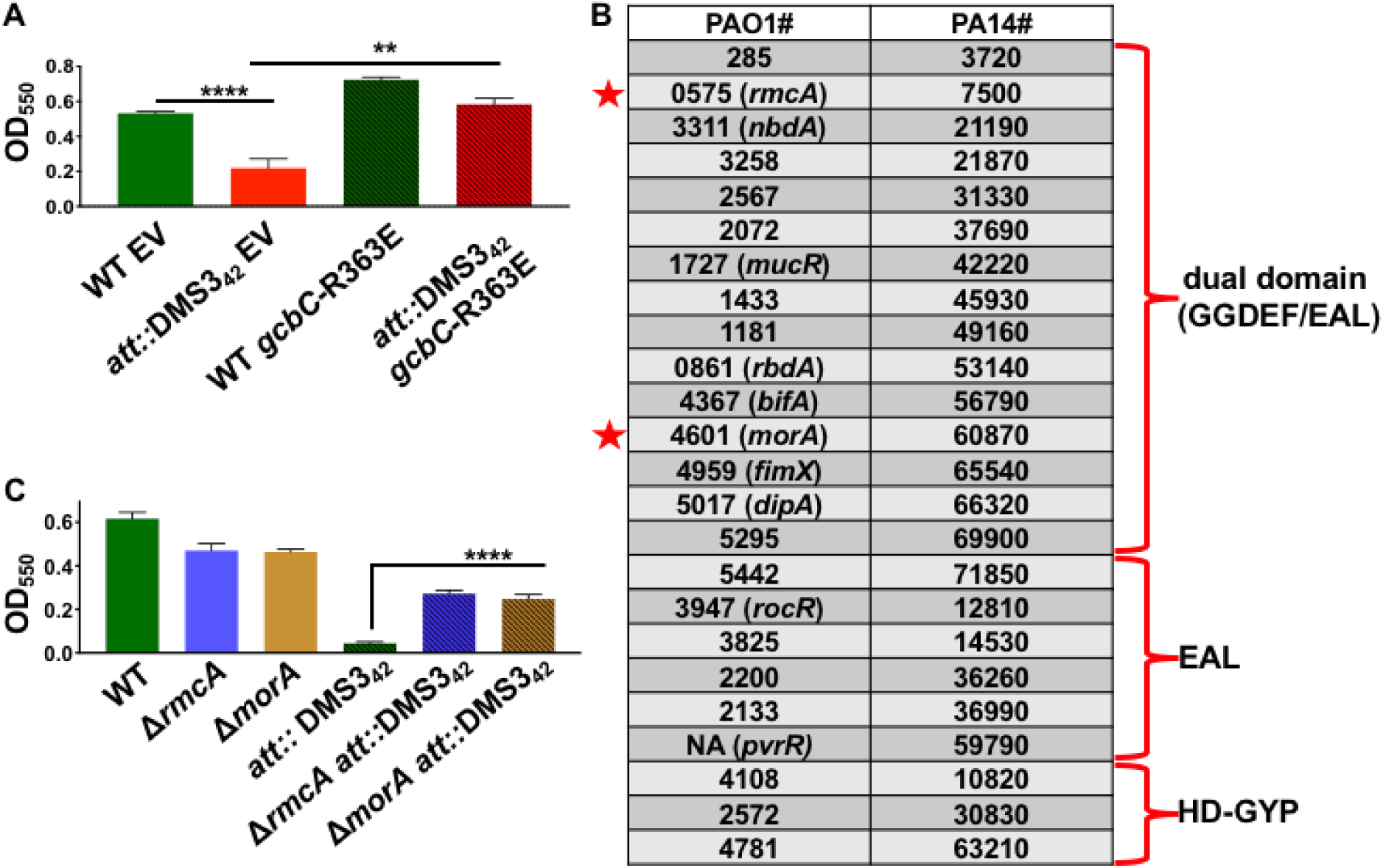
CRISPR-activated *P. aeruginosa* is unable to maintain a biofilm but can be rescued by modulating c-di-GMP levels. (A) Biofilm assays were performed as detailed in Materials and Methods and grown in M63 medium supplemented with 0.4% L-arginine, which we refer to as the “biofilm medium” in the text. WT (*P. aeruginosa* PA14) and the strain carrying the DMS3_42_ insertion at the *att* site (*att*::DMS3_42_), carrying either the empty vector (EV) or a plasmid expressing the constitutively-active diguanylate cyclase GcbC*-*R363E, were grown statically for 12 h then biofilm formation assessed. The OD_550_ value represents a measure of the biofilm formed. Error bars represent standard deviation of the results from three biological replicates each performed with three technical replicates. *** indicates a difference in biofilm levels that is significantly different at a P value of <0.001 for the indicated strains. (B) Previous work by Ha and colleagues (24) created in-frame, unmarked deletions of all putative phosphodiesterases that were identified via SMART analyses of the *P. aeruginosa* genome. The deletion constructs were introduced into the *att*::DMS3_42_ background, and mutations in two genes indicated by red stars, *rmcA* (Δ*rmcA att*::DMS3_42_) and Δ*morA* (Δ*morA att*::DMS3_42_) were found to produce significantly more biofilm compared to the *att*::DMS3_42_ strain. (C) The Δ*rmcA* and Δ*morA* mutations in the WT or the CRISPR-activated background (*att*::DMS3_42_) were assessed for biofilm formation after 12h. The OD_550_ value represents a measure of the biofilm formed. Error bars represent standard deviation of the results from three biological replicates each performed with three technical replicates. **** indicates a difference in biofilm levels that is significantly different at a P value <0.0001 compared to the respective WT *att*::DMS3_42_ strain.

Given that the biofilm defect of the *att*::DMS3_42_ strain could be rescued by heterologously expressing a c-di-GMP-synthesizing enzyme, we reasoned that the same effect might/should be observed by mutating one or more genes encoding PDEs. To this end, we built on work from Ha *et. al*. who used the SMART algorithm to analyze the genome of *P. aeruginosa* for encoded proteins with motifs related to c-di-GMP metabolism (23, 24). This approach identified 24 candidate PDEs with either EAL domains alone (6), dual GGDEF/EAL domains (15), or HD-GYP domains (3) (**Fig. 1B**). Individual in-frame deletions for all these PDEs were constructed in the *att*::DMS3_42_ background and these strains were then assayed for biofilm formation. Of all the deletions of genes coding for PDE that were constructed, only two of them, Δ*rmcA* and Δ*morA*, exhibited a significant rescue of biofilm biomass at 12 h compared to the parental *att*::DMS3_42_ background (**Fig. 1C**). While both *rmcA* and *morA* encode for dual (GGDEF and EAL) domains, previous work suggests that these proteins behave predominately as PDEs (25, 26). These data suggest that the defect observed in the *att*::DMS3_42_ strain can be rescued by increasing concentrations of c-di-GMP either via heterologous expression of the GcbC-R63E protein (**Fig. 1A**) or through the loss of either of the two PDEs encoded by the *rmcA* and *morA* genes (**Fig. 1C**). The observation that none of the other PDEs tested could rescue the biofilm formation defect of the *att*::DMS3_42_ strain (**Fig. 1B**) suggests a specific role for these particular PDEs, a point we address below.

### The Δ*rmcA* or Δ*morA* mutants are defective in the later stages of biofilm formation in a static assay

We next investigated the biofilm phenotypes of the Δ*rmcA* and Δ*morA* mutants in a WT background (no DMS3_42_-mediated CRISPR activation) using a 96-well dish static biofilm assay over a 48 h window. These mutants were able to form a biofilm similar to the WT over the first ∼12 h of the assay (**Fig. 2A**). Between 12 and 24 h, while WT biofilm biomass continues to increase, the biomass of the Δ*rmcA* and Δ*morA* mutants plateau and begin to decrease, and increasingly exhibit a defect in biofilm biomass out to 48 hr. Between 24 and 48 h the WT biofilm is maintained with no obvious loss of biomass in this assay.

**Figure 2.**
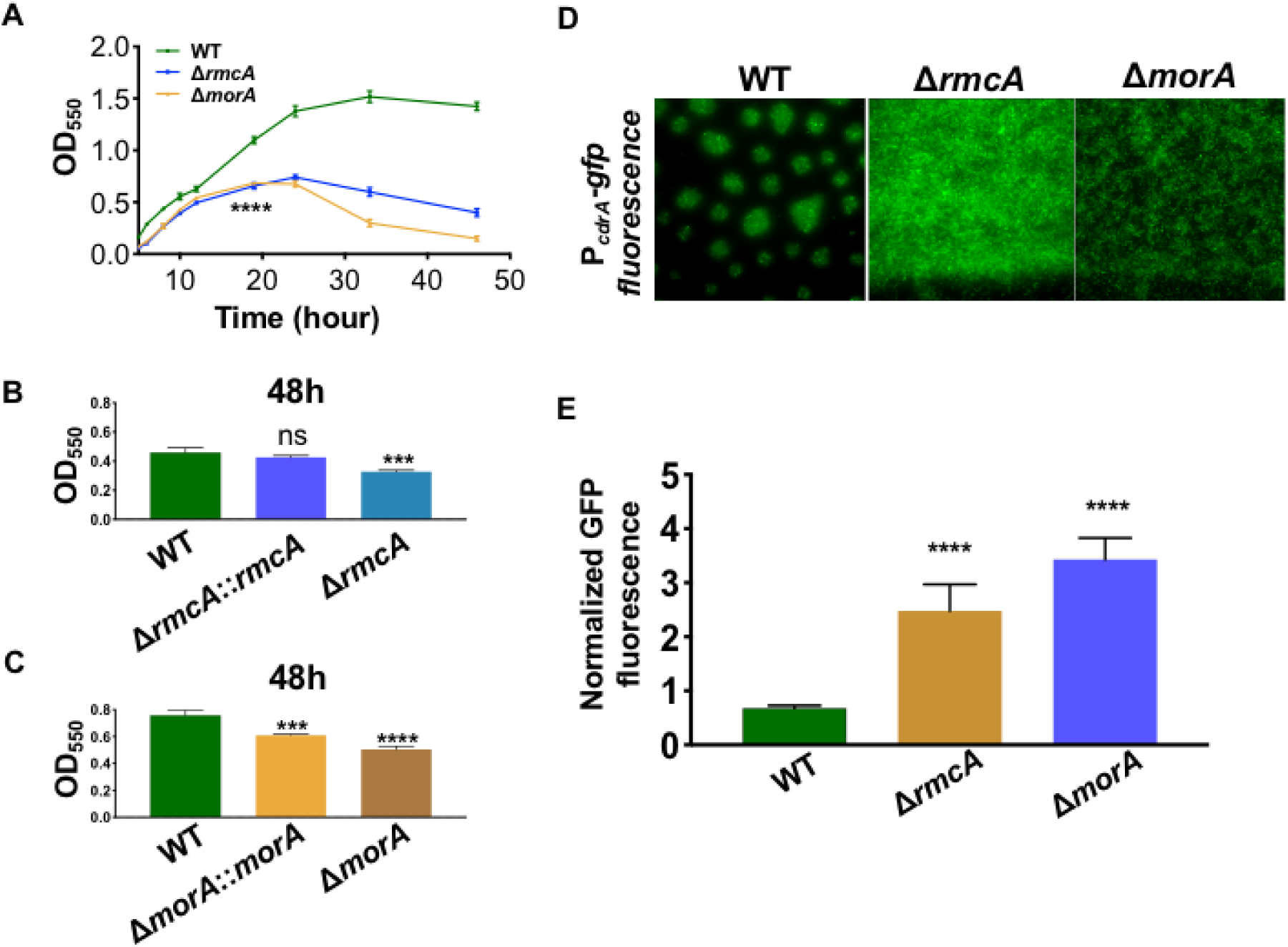
The Δ*rmcA* and Δ*morA* mutants are defective in biofilm maintenance and show increased levels of c-di-GMP. (A) The WT strain (*P. aeruginosa* PA14) and the Δ*rmcA* and Δ*morA* mutant strains were grown statically in a 96 well dish assay using M63 medium supplemented with 0.4% L-arginine, and the biofilm formed measured at the indicated timepoints. *** indicates a difference in biofilm that is significantly different at a P value of <0.001 at the indicated time point and each subsequent time point. (B) The WT, Δ*rmcA* mutant, and the Δ*rmcA*::*rmcA*^*+*^ complemented strain were grown for 48 h prior to crystal violet staining to assess the extent of biofilm formation. *** indicates a difference that is significantly different at a P value of <0.01 compared to the WT; ns is not significant. (C) The WT, Δ*morA* mutant, and the Δ*morA*::*morA*^*+*^ complemented strain were grown for 48 h prior to crystal violet staining to assess the extent of biofilm formation. *** indicates a difference that is significantly different at a P value of <0.01 compared to the WT. (D) The WT and the Δ*rmcA* and Δ*morA* mutants carrying the P_*cdrA*_-*gfp* fusion expressed from a multicopy plasmid were grown along the air-liquid interface of 18 mm glass coverslips and were imaged after 24 hr. Fluorescent microscopy was used to determine GFP signal intensity. (E) Quantification of GFP signal intensity for the strains described in panel D. Fluorescent microscopy was used to determine GFP signal intensity as a measure of c-di-GMP production, which was normalized to constitutively active mKO fluorescent protein. **** indicates a difference in biofilm that is significantly different at a P value < 0.0001 compared to the WT strain. In all panels, error bars represent standard deviation from three biological replicates each performed with three technical replicates.

To assess if the behavior of Δ*rmcA* and Δ*morA* is dependent on the type of carbon source provided, we replaced the carbon source used in our standard medium, L-arginine, with pyruvate. We found that that the Δ*rmcA* and Δ*morA* mutant biofilms also exhibit a late-stage defect at 36 h in this medium, while the 12 h biofilms were similar to WT (**Fig. S1**). To verify that the loss of biofilm that we observed in late timepoints of our kinetic assay was dependent on the absence of either RmcA or MorA we complemented these mutants, and found that the late stage biofilm defect observed in these mutants could be rescued by a wild-type copy of *rmcA* and to a lesser extent for *morA*, respectively (**Fig. 2B & C**).

### The Δ*rmcA* and Δ*morA* mutants have increased c-di-GMP levels

Given that RmcA and MorA are predicted PDEs (25, 26), we would expect that the Δ*rmcA* and Δ*morA* mutants would have increased levels of c-di-GMP compared to the WT strain. To test this hypothesis, we measured total c-di-GMP in the mutants versus WT using a P_*cdrA*_-*gfp* promoter fusion to assess transcriptional activity of *cdrA*, a gene that is positively regulated by c-di-GMP (27). The GFP signal serves as a surrogate for c-di-GMP levels. We normalized the amount of GFP signal produced to a constitutively expressed orange fluorescent protein, mKO (28), integrated into the *att* chromosomal insertion site. This assay provided us with an indirect, but normalized measure of the relative c-di-GMP concentration in WT versus mutant backgrounds. Biofilms were grown in a static assay on coverslips that were partially submerged in buffered biofilm medium containing 0.4% arginine in a 12-well dish, as described (29). The coverslips were then incubated for 24 h, similar to the static biofilm assays shown in **Figure 1** and **Figure 2**, prior to imaging of the air-liquid interface (ALI) with brightfield and fluorescence microscopy.

Mutation of either *rmcA* or *morA* resulted in an increase of P_*cdrA*_-*gfp* fluorescence (**Fig. 2D**). The structures of the WT and mutant biofilms also differed, with the WT generating typical “mushroom-like” colonies, while the mutants formed a uniform layer of cells. Thus, it was important to examine the normalized signal intensity of individual cells to accurately compare the average signal intensity between strains. On a cell-by-cell basis, with values normalized to mKO signal intensity, both the Δ*rmcA* and Δ*morA* mutants exhibited a significant increase in signal intensity from the P_*cdrA*_-*gfp* reporter of 3.6- and 5.0-fold over WT, respectively (**Fig. 2E**). These results are consistent with the previous reports that both *rmcA* and *morA* encode for active PDEs and that the loss of these enzymes would result in reduced degradation of c-di-GMP.

Given the observation that strain backgrounds with a late-stage biofilm deficiency also had elevated c-di-GMP, we hypothesized that this deficiency could be induced in WT or enhanced in Δ*rmcA* and Δ*morA* mutants simply by increasing c-di-GMP production. To test this hypothesis, we measured biofilm levels at several time points in WT and mutant backgrounds that were heterologously expressing *gcbC*-R363E from a plasmid. As exhibited in **Fig. S2** however, increasing c-di-GMP production alone is insufficient to induce a late-stage biofilm deficiency in the WT, nor is it capable of accelerating the kinetics of the observed biofilm defect in the Δ*rmcA* and Δ*morA* mutant backgrounds. These data suggest that the observed phenotypes are linked to the loss of *rmcA* and *morA* specifically, rather than to a general increase in c-di-GMP levels.

### The inability of Δ*rmcA* and Δ*morA* to maintain a late-stage biofilm is correlated to nutrient limitation

The death observed in late stage biofilms in the static assay indicated that perhaps, as nutrients become limited in this batch culture, the Δ*rmcA* and Δ*morA* mutants were unable to adapt to these nutrient-limited conditions and thus exhibited reduced viability. We tested this hypothesis in two ways. First, we conducted static 96-well biofilm assays in which biofilm formation was measured early (12 h) and late (48 h), and then compared these results to biofilms grown for the same 48 h, but with periodic (every 12 h) removal of spent medium followed by the addition of fresh medium. That is, periodic replacement of the medium provided cells with regular access to fresh nutrients even in the batch culture system. We found that, compared to late timepoints in the standard assay (no medium replacement) wherein the Δ*rmcA* and Δ*morA* mutants exhibited a biofilm defect (55.7% and 82.5% reduction, respectively; **Fig. S3A**), the removal and replacement of spent medium with fresh medium results in the reduction of the magnitude of this defect to an 11.6% reduction in Δ*rmcA* and an 33.1% reduction in Δ*morA* (**Fig. S3B**), levels comparable to that observed for an early stage biofilm formed by the Δ*morA* mutant at 12 h (**Fig. S3C**).

We next assessed whether oxidative stress could result in a similar cell death phenotype. We grew biofilms for 12 h in a static 96 well assay, a time point wherein we have detected minimal evidence of starvation responses within the biofilm (**Fig. 2A and S4A**). We next replaced the spent medium with fresh medium supplemented either with 20 mM H_2_O_2_ (**Fig. S4B**) or no H_2_O_2_ (**Fig. S4C**) and then incubated the biofilms for an additional 6 h. The addition of H_2_O_2_ had no impact on the WT or mutant biofilms (**Fig. S4B**) compared to the control (**Fig. S4C**). Taken together, these results are consistent with the hypothesis that the defect that we observe in the Δ*rmcA* and Δ*morA* mutants is due to an inability to appropriately respond to nutrient limitation, rather than a general stress response.

### The inability of the Δ*rmcA* and Δ*morA* mutants to maintain biofilm correlates with increased cell death

To explain the loss of late-stage biofilms in the Δ*rmcA* and Δ*morA* mutants we tested the hypothesis that these mutants were dying, thereby causing the loss of biofilm biomass. We imaged biofilms after staining with the LIVE/DEAD *Bac*Light kit (Molecular Probes). This assay allowed us to determine the ratio of cells that are viable (i.e., those stained green by membrane-permeable Syto9) to those that are dead (i.e., cells with compromised membranes that are stained red by membrane-impermeable propidium iodide). Biofilms were stained with *Bac*Light after 16 h or 48 h of static growth in the ALI assay in 12 well plates, as described above, and the data plotted as the ratio of live cells (green) to dead cells (red).

After 16h, biofilms of all strain backgrounds were comprised of predominantly viable cells (**Fig. 3A**, top row) with the Δ*morA* mutant displaying a live/dead ratio similar to WT, while the Δ*rmcA* mutant live/dead ratio was significantly higher than the WT even at this early time point (**Fig. 3B**, left panel). After 48 h in the static assay, however, the Δ*rmcA* and Δ*morA* mutants were significantly less viable than WT (**Fig. 3A**, bottom row) with the live/dead ratios of Δ*rmcA* and Δ*morA* mutants reduced by 55.7% and 43.3%, respectively (**Fig. 3B**, right panel). While both mutants exhibited a reduced ratio of live/dead cells, individual comparisons of Syto9 and PI (**Fig. S5A and B**, respectively) reveal that the reduction observed in Δ*rmcA* mutant is driven both by an increase in the number of dead cells and a decrease in live cells, whereas while in Δ*morA* mutant the change in this ratio reflects primarily the loss of viable (green) cells. The inability to detect a significant increase in dead cells within late-stage Δ*morA* biofilm could be due to an earlier onset of death followed by the sloughing off of dead cells prior to microscopy at 48 h, a conclusion consistent with the findings presented below.

**Figure 3.**
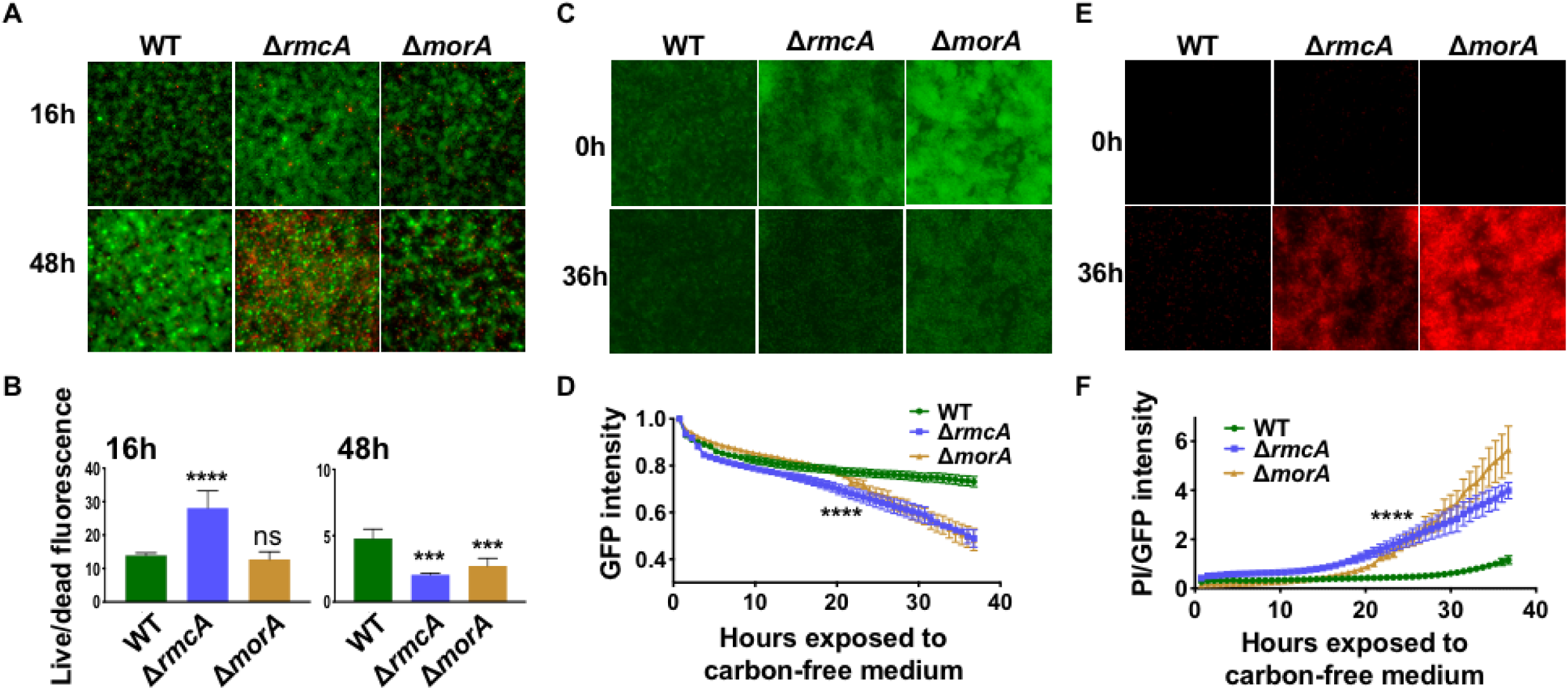
The biofilm defect of Δ*rmcA* and Δ*morA* mutants coincides with cell death. The WT strain (*P. aeruginosa* PA14) and the Δ*rmcA* and Δ*morA* mutants were grown along the air-liquid interface of 18 mm glass coverslips. At 24 h the coverslips were removed, washed in PBS and stained with Syto-9 and propidium iodide (PI). Fluorescent microscopy was used to measure the Syto-9 and PI fluorescence (panel A) and image intensity assessed as a ratio live (Syto-9) to dead (PI) was determined at 16 h (panel B, left) and 48 h (panel B, right). Error bars represent standard deviation from three biological replicates each performed with three technical replicates. *** and **** indicate a difference in biofilm that is significantly different at a P value of <0.001 and <0.0001, respectively, compared to the WT; ns, not significant. (C) The WT strain and the Δ*rmcA* and Δ*morA* mutants carrying plasmid pSMC21 were grown for 23 h in biofilm medium containing 0.4% L-arginine and stained under flow for an additional 1 h in biofilm medium containing propidium iodide (PI). After this 24 h period of incubation, the biofilm medium was replaced with medium containing PI and lacking arginine; this was considered to be time = 0 h of nutrient limitation (top panel). The biofilms imaged after 36 h of nutrient limitation are shown at the bottom of the panel. (D) GFP fluorescence of the biofilms in panel C were measured every 45 minutes for 36 hrs. WT and the mutants were assessed for changes in GFP fluorescence and plotted as a fraction of GFP signal at the start of nutrient limitation. (E) PI staining of the corresponding biofilms from panel C are shown just prior to nutrient limitation and after 36 h of nutrient limitation. (F) PI straining normalized to GFP fluorescence of pSMC21 is plotted for data acquired in panels C and E every 45 minutes. Error bars represent standard deviation from three biological replicates each performed with three technical replicates. **** indicates a difference in biofilm that is significantly different at a P value of <0.0001, a level of significance observed after 23 h of exposure to nutrient-limited conditions, and at all subsequent time points.

To address whether we could recapitulate the impact of nutrient-limited conditions observed in our static assays, we utilized a microfluidic device that allowed us to observe biofilm dynamics in real time and to manipulate the amount of nutrients provided. We first confirmed that all three strains could form biofilm within a microfluidic chamber. To monitor biofilm formation, we introduced the pSMC21 plasmid, which constitutively expresses GFP (30, 31), into each of the strains. The bacteria were inoculated into the microfluidic chamber, allowed to attach for 1 h prior to the start of flow (0.5 μl/min), then monitored at 45 min intervals using fluorescence microscopy over the first 12 h of biofilm formation. The Δ*rmcA* and Δ*morA* mutants are able to form robust biofilms in the microfluidic chamber (**Fig. S6**).

We hypothesized that we could recapitulate in a microfluidic chamber the nutrient-limited conditions that developed over late timepoints in the static assays by establishing the biofilms in a medium that contained a carbon source, then irrigating the biofilms with medium lacking a carbon source. To test this idea, we allowed biofilms of the WT and mutants carrying the GFP-expressing plasmid pSMC21 to form a biofilm for 24 h in biofilm medium with arginine as the carbon source, then switched to biofilm medium lacking arginine. One hour before the switch to nutrient-limited conditions, we stained the microfluidic chamber-grown biofilms with PI to label non-viable cells and thus establish a baseline of non-viable cells before inducing nutrient limitation. As shown in **Figure 3C** and **Figure 3E**, at time zero before nutrient limitation is induced, all three strains formed a robust biofilm and showed minimal non-viable cells.

To assess if nutrient limitation differentially impacted loss of biofilm in the WT compared to the mutants, we normalized the GFP signal to the start of nutrient limitation for each strain (t = 0 in **Fig. 3C**, top), then recorded the change in GFP intensity over the subsequent 36 h of exposure to carbon-free medium. While WT was largely able to maintain the biofilm over the course of the assay, the lack of carbon in the growth medium accelerated the loss of biomass in both the Δ*rmcA* and Δ*morA* mutant biofilms. This was evidenced by the greater reduction of the GFP-mediated signal intensity in the mutants compared to the WT during the 36 h period of carbon limitation (**Fig. 3C-D**). The WT lost ∼25% of biomass compared to a >50% reduction for both the Δ*rmcA* and Δ*morA* mutants over the 36 h of the experiment.

To assess if these mutants lost biofilm biomass over the course of nutrient limitation due to increased death, as observed in the static assay in **Figure 3A**, we measured the intensity of PI over time and found that both the Δ*rmcA* and Δ*morA* mutants exhibited rapidly increasing PI staining in absolute terms (**Fig. 3E**) and as a ratio of GFP fluorescent intensity, which served as a measure of total biofilm biomass of viable cells (**Fig. 3F**). These data provide further evidence that both the Δ*rmcA* and Δ*morA* mutants are susceptible to nutrient-limited conditions when grown in a biofilm, and that loss of late-stage biofilm biomass coincides with cell death.

To determine if dispersal of the biofilm could contribute to the observed loss of biofilm biomass upon nutrient limitation, we measured the viable count (CFU) of bacteria dispersing from biofilms within a microfluidic device prior to and after nutrient limitation. As shown in **Figure S7**, effluent-derived cells of all strains were similar 2 h prior to, as well 12 and 18 h after, the switch to arginine-free medium. Only in the final timepoint (24 h) did the viable count of the Δ*rmcA* and Δ*morA* effluent increase compared to WT. These data suggest that cell death, and not dispersal, is the primary driver of the loss of biomass observed in these PDE mutants during nutrient limitation.

### A stringent response mutant phenocopies the biofilm cell death of Δ*rmcA* and Δ*morA* mutants during nutrient limitation

The ability of the Δ*rmcA* and Δ*morA* mutants to maintain a biofilm when nutrients were present coupled with the decrease in viability during nutrient limitation suggested that these mutants were unable to mediate the appropriate responses needed for persistence when resources become limiting. This conclusion was further supported by the observation that the biofilm defect in the static assay for the Δ*rmcA* and Δ*morA* mutants could be rescued simply by adding fresh medium (**Fig. S3**). Based on these data, we hypothesized that loss of RmcA or MorA function results in the inability to appropriately navigate nutrient limited conditions.

If this hypothesis is correct, other strains defective in the nutrient limitation-response should have a similar phenotype. To test this prediction, we assessed the phenotype of a Δ*relA* Δ*spoT* double mutant, which is unable to either make or degrade the alarmone (p)ppGpp critical for the stringent response, for its ability to respond to nutrient limitation when grown in a biofilm. We first assessed biofilm formation of the Δ*relA* Δ*spoT* mutant in static assays and found that, like the Δ*rmcA* and Δ*morA* mutants (**Fig. 2A**), the Δ*relA* Δ*spoT* mutant could form a biofilm (albeit at a level lower than the WT) at early time points and biofilm biomass was reduced at later time points (**Fig. 4A**).

**Figure 4.**
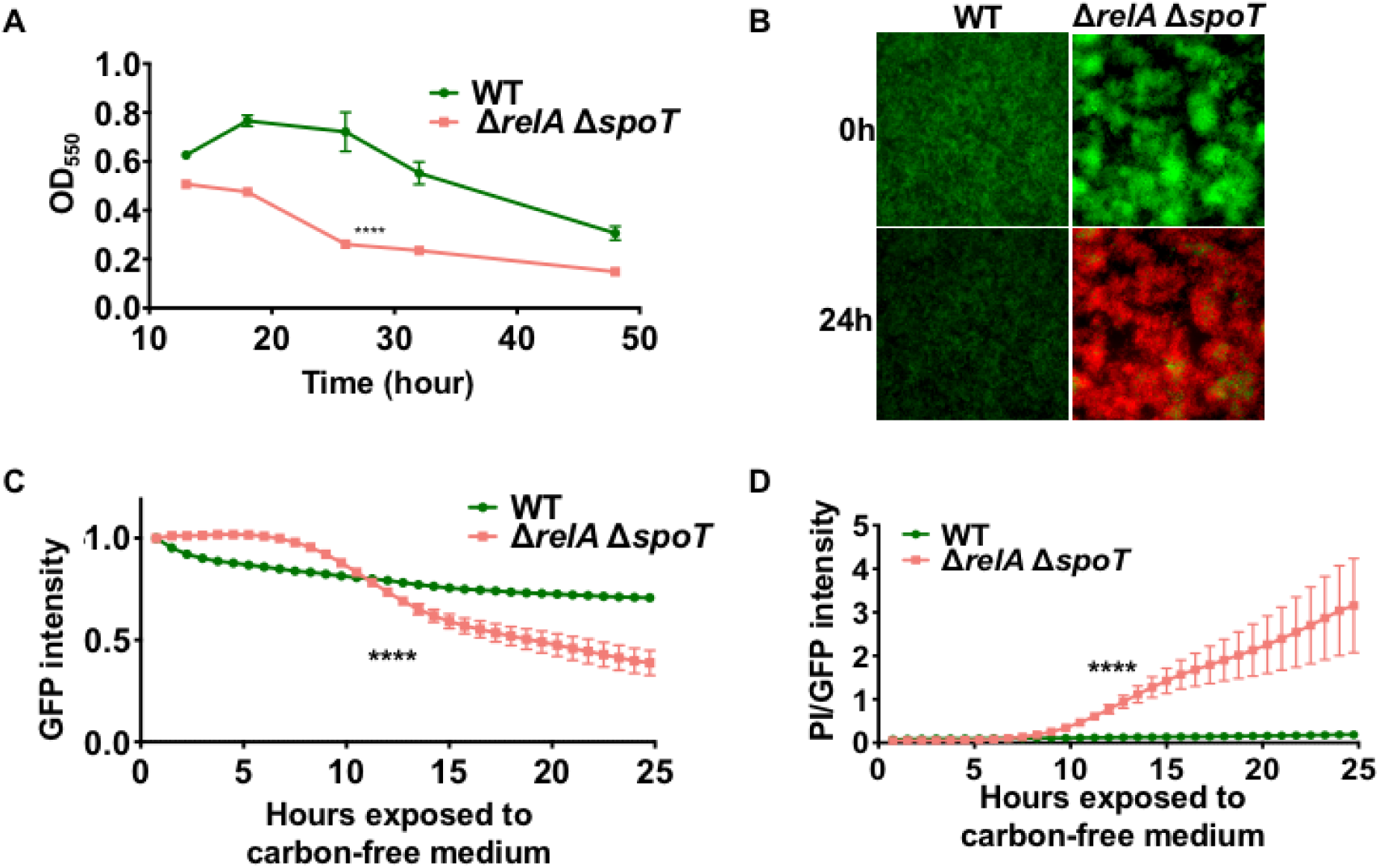
Loss of stringent response phenocopies biofilm defect and cell death observed for the PDE mutants. (A) The WT (*P. aeruginosa* PA14) strain and the Δ*relA*Δ*spoT* double mutant strain were grown statically in a 96 well dish assay using M63 medium supplemented with 0.4% L-arginine, and biofilm formed was measured at the indicated timepoints. (B) The WT strain and the Δ*relA*Δ*spoT* double mutant, both carrying pSMC21, were grown for 23 h using a microfluidic device in biofilm medium containing 0.4% L-arginine and stained under flow for an additional 1 h in biofilm medium containing propidium iodide (PI). The biofilm medium was then replaced with medium containing PI and lacking arginine. The fluorescence due to GFP and PI for the WT strain and the Δ*relA*Δ*spoT* double mutant are shown at the start of nutrient limitation (0 hr, top) and after 24 h (bottom) in the microfluidic chamber. (C) GFP fluorescence was measured every 45 minutes for 24 h after the initiation of nutrient limitation. The WT strain and the Δ*relA*Δ*spoT* double mutant strain were assessed for changes in GFP fluorescence, which was normalized to the GFP signal at the start of nutrient limitation, which is set to 1. (D) The ratio of PI to GFP during ∼25 h of nutrient limitation in the microfluidic chamber is presented as a measure of cell viability, with larger values indicating more cell death. For all panels, error bars represent standard deviation from three biological replicates each performed with three technical replicates. Shown is a representative experiment. **** indicates a difference in biofilm that is significantly different at a P value of <0.0001 at the indicated time point and each subsequent time point.

Next we used a microfluidic chamber where nutrient-limited conditions could be induced via introduction of carbon-free medium. Using the same conditions we used above (**Fig. 3**), we found that for the Δ*relA* Δ*spoT* mutant, which is unable to mount a stringent response, the loss of biofilm biomass (**Fig. 4B**) and viability (**Fig. 4C-D**) occurs concurrently with nutrient limitation.

### Loss of RmcA and MorA function is associated with increased Pel polysaccharide production in the biofilm

Increased c-di-GMP is typically associated with enhanced biofilm formation. To reconcile how mutants that have a late-stage biofilm defect (**Fig. 3**) also produce increased c-di-GMP (**Fig. 2C-D**), we hypothesized that the loss of biofilm observed in the Δ*rmcA* and Δ*morA* mutants was the result of untimely cellular investment in energetically expensive products. Such a view is consistent with the kinetics of the defect observed for the Δ*rmcA* and Δ*morA* mutants, which becomes increasingly evident after ∼30 h of growth in static conditions, when nutrients are likely depleted, and after a shift to carbon-free medium in the microfluidic device.

To evaluate whether the increased concentration of c-di-GMP in late-stage biofilms also resulted in altered phenotypes relevant to biofilm formation, we assessed production of extracellular polysaccharide (EPS) in the WT and the Δ*rmcA* and Δ*morA* mutants. In *P. aeruginosa* PA14 the dinucleotide c-di-GMP up-regulates Pel production. Both mutants showed enhanced pellicle production compared to the WT, with accumulated biomass on the tubes of overnight-grown planktonic cultures (**Fig. 5A**). We also employed Congo Red (CR), a dye which can be used as a qualitative indicator of the presence of EPS, combined with colony biofilm assays on agar medium. The Δ*rmcA* and Δ*morA* mutants showed enhanced CR binding after 4-5 days compared to the WT (**Fig. 5B**), consistent with the view that the loss of RmcA and MorA results in increased c-di-GMP and EPS production.

**Figure 5.**
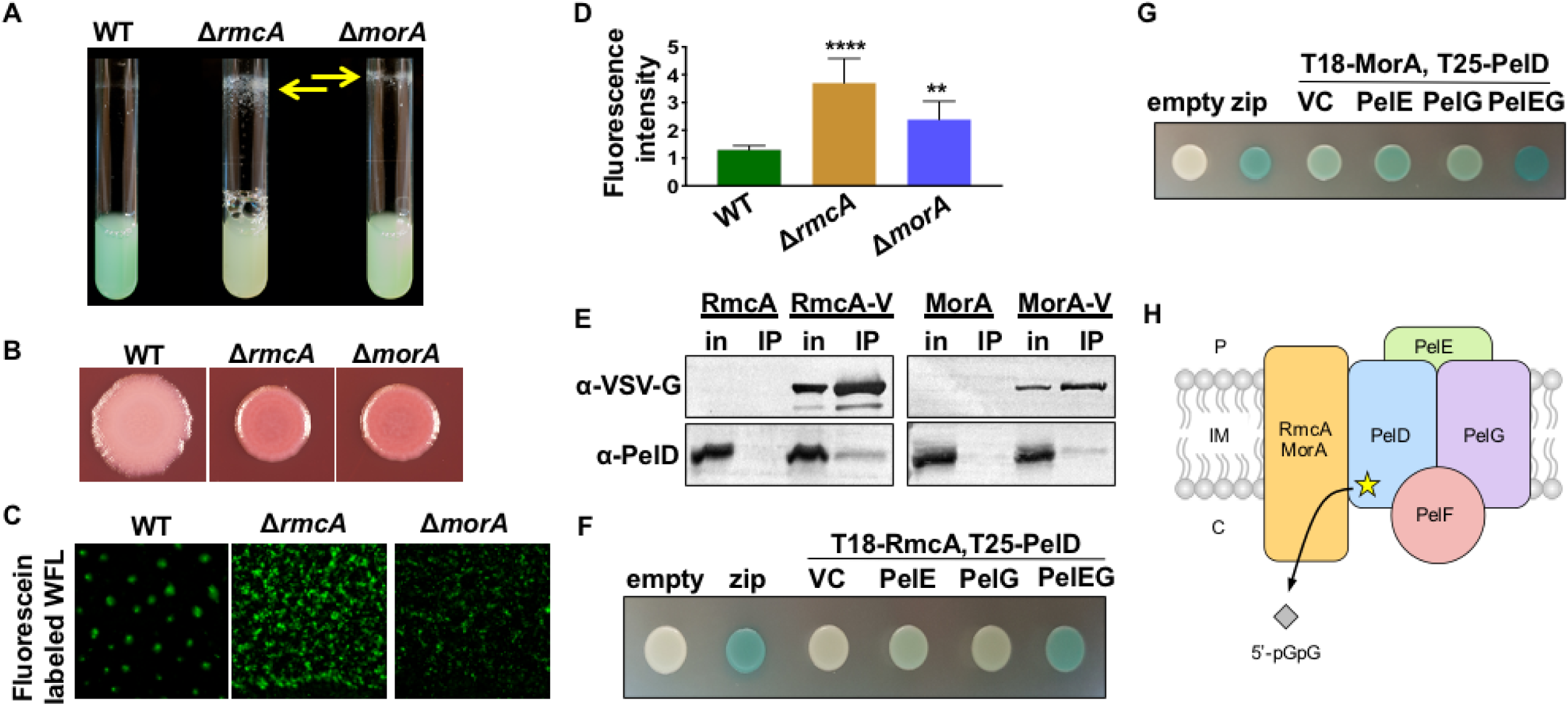
The loss of RmcA or MorA function results in increased Pel production likely through physical interaction with the Pel complex. (A) Cultures of the indicated *P. aeruginosa* PA14 strains were inoculated into lysogeny broth (LB) and imaged after overnight growth at 37°C. The resulting wall-associated material is indicated by the yellow arrows. (B) Congo Red plate assays of the indicated *P. aeruginosa* PA14 strains are shown. The plates were incubated for 24 h at 37°C then at room temperature for an additional 4 days. (C) The WT (*P. aeruginosa* PA14) strain and the Δ*rmcA* and Δ*morA* mutants were grown as a biofilm in the ALI assay for 18 h in medium containing fluorescein-labeled WGA and washed before imaging by fluorescence microscopy. (D) Quantification of the experiment performed in panel C. The fluorescence attributable to the fluorescein-labeled WGA was normalized to the mKO fluorescent protein expressed from the chromosome of each strain and plotted. Error bars represent standard deviation from three biological replicates each performed with three technical replicates. ** and **** indicate a difference in biofilm that is significantly different at a P value of <0.01 and 0.0001, respectively. (E) Co-IP from solubilized *P. aeruginosa* PAO1 inner membranes expressing VSV-G-tagged RmcA (RmcA-V, *left*) or MorA (MorA-V, *right*) as the bait. The corresponding untagged proteins (RmcA, MorA) were used as a negative binding control. Proteins applied to the α-VSV-G co-IP resin (in, input) and the elution from the resin after washing (IP, immunoprecipitated) were analyzed by Western blot using VSV-G and PelD specific antibodies, as indicated. (F) Representative colony images for the analysis of interactions between RmcA, fused at the N-terminus to the T18 domain of *Bordetella pertussis* adenylate cyclase toxin (T18-RmcA), and PelD fused to the T25 adenylate cyclase domain at the N-terminus (T25-PelD), by BACTH using solid media containing X-Gal as a reporter. The assay was modified through the additional of an empty vector control (VC) for expression of untagged PelE, untagged PelG, or both PelE and PelG (PelEG). A blue colony indicates a positive result in this assay. Empty vectors expressing the T18 or T25 domain alone (empty) were used as a negative control. Fusion of the T18 and T25 domains to a leucine zipper motif (zip) was used as a positive control. (G) Representative colony images for the analysis of interactions between MorA, fused to the T18 domain at the N-terminus (T18-RmcA), and T25-PelD by BACTH. Experiments were performed as described for panel F. Note: The proteins used in these BATCH assays were derived from *P. aeruginosa* PAO1, which are 99% identical to the *P. aeruginosa* PA14 proteins. (H) A model for the RmcA and MorA proteins in complex with the Pel biosynthetic complex (PelDEFG). 5’-pGpG (diamonds) represents the product of the PDE-mediated degradation of c-di-GMP.

To test if the Pel polysaccharide specifically was being overproduced in the Δ*rmcA* and Δ*morA* mutants, we used fluorescein-labelled *Wisteria floribunda* lectin (WFL) which binds preferentially to carbohydrate structures that terminate in *N*-acetylgalactosamine. This lectin has been shown to bind specifically to the Pel polysaccharide (32). WFL was used to stain statically grown ALI biofilms and the fluorescent signal was normalized to a constitutively expressed fluorescent tag, mKO, as described above. Biofilms of the Δ*rmcA* and Δ*morA* mutants demonstrated elevated WFL binding (**Fig. 5C**), with signals from the Δ*rmcA* and Δ*morA* mutants significantly increased by 2.8- and 1.9-fold above WT, respectively (**Fig. 5D**). Combined, these data suggest that both the Δ*rmcA* and Δ*morA* mutants produce increased levels of the Pel EPS when growing as a biofilm, consistent with the elevated levels of c-di-GMP observed in these mutants (**Fig. 2**).

To further probe whether high levels of Pel expression in the Δ*rmcA* and Δ*morA* mutants was necessary and/or sufficient to induce late cell death. First, we introduced a Δ*pelA* mutation into the Δ*rmcA* and Δ*morA* backgrounds, but unfortunately, the Δ*rmcA* Δ*pelA* and Δ*morA* Δ*pelA* double mutants were defective for establishing a biofilm (**Fig. S8A**), which is expected given the role of Pel in biofilm formation (40), so we could not perform the desired analyses in these strains. Instead, we used a strain that allowed us to artificially induce Pel expression to high levels – this strain did not show enhanced death in late stage biofilms (**Fig. S8B**). This second observation indicates that while Pel production may contribute to late stage cell death in the Δ*rmcA* and Δ*morA* mutants, it is not sufficient to drive this phenotype. That is, the high levels of c-di-GMP in the Δ*rmcA* and Δ*morA* mutants may have additional negative impacts on the cell in the context of a nutritionally limited, mature biofilm.

### RmcA and MorA physically interact with PelD

The data presented thus far suggests that appropriate Pel regulation is lost when cells lack RmcA or MorA function. The biosynthesis of Pel is regulated at the transcriptional and post-translational levels by the c-di-GMP-binding effector proteins FleQ and PelD, respectively (33, 34). DGCs and PDEs can influence the activation state of effector proteins through the alteration of global intracellular c-di-GMP pools or via specific interaction with effectors via local signaling events (35). RmcA, MorA, and PelD localize to the inner membrane due to the presence of one or more predicted transmembrane helices (25, 36, 37), therefore we hypothesized that RmcA and/or MorA may influence Pel biosynthesis through direct interactions with PelD.

To assess possible RmcA/MorA/PelD interactions, a vesicular stomatitis virus glycoprotein (VSV-G)-tag was added to the C-terminus of RmcA and MorA (RmcA-V and MorA-V), and the genes expressing these tagged proteins were integrated at the neutral chromosomal *att*::Tn*7* site under the control of the *araC*-P_BAD_ promoter in *P. aeruginosa* strains lacking a native copy of the *rmcA* or *morA* genes, respectively. To ensure that the activity of RmcA or MorA did not influence the c-di-GMP-dependent transcription of the *pel* operon by FleQ, the Pel overproducing strain *P. aeruginosa* PAO1 Δ*wspF* Δ*psl* P_BAD_*pel* was utilized (38). Co-immunoprecipitation (co-IP) was performed from solubilized, enriched inner membranes of *P. aeruginosa* overexpressing the RmcA-V or MorA-V tagged proteins and the protein components encoded by the *pel* operon by the addition of L-arabinose to culture media. In each experiment, PelD was detected in the eluate via Western blot when RmcA-V or MorA-V were supplied as the bait, but not in the untagged control eluate (**Fig. 5E**). These data suggest that RmcA and MorA may interact with PelD to exert their control over Pel biosynthesis.

While the data gathered by co-IP suggests an interaction between RmcA/MorA and PelD, it does not distinguish between direct interactions or those mediated by other unknown proteins. To validate these findings using a different approach, interactions between RmcA/MorA and PelD were analyzed using bacterial two hybrid (BACTH) assays. In these experiments, the inactive T18 fragment of *Bordetella pertussis* adenylate cyclase toxin was fused to the N-terminus of RmcA or MorA (T18-RmcA or T18-MorA), while the inactive T25 adenylate cyclase fragment was fused to the N-terminus of PelD (T25-PelD). Interaction between T18-RmcA/T18-MorA and T25-PelD would reconstitute adenylate cyclase enzymatic activity, and lead to the production of blue colonies when analyzed in the *E. coli* BTH101 reporter strain grown on agar medium containing X-Gal.

When interactions between T18-RmcA and T25-PelD were examined in the BACTH assay, white colonies were observed, indicative of a negative result (**Fig. 5F**). Since interactions between RmcA and PelD were identified by co-IP in a *P. aeruginosa* background where the entire *pel* operon was overexpressed (**Fig. 5E**), and PelD directly interacts with both PelE and PelG to form the inner membrane Pel synthase complex regardless of its c-di-GMP binding capability (36), we reasoned that expressing untagged PelE, PelG, or both co-polymerase proteins alongside T18-RmcA and T25-PelD would better imitate the physiological conditions under which this interaction is presumed to occur. When these modified BACTH experiments were performed, we observed bright blue colonies comparable to the positive control when both PelE and PelG were co-expressed with T18-RmcA and T25-PelD, but only very faint blue to white colonies when PelE or PelG were singly co-expressed (**Fig. 5F**). Similar results were obtained when the modified BACTH experiment was performed with T18-MorA and T25-PelD, where a deep blue colony indicative of a positive result was observed when both untagged PelE and PelG were co-expressed (**Fig. 5G**). However, unlike with RmcA, a weak-to-moderate positive result was also obtained when only untagged PelE, untagged PelG, or even an empty vector control was present (**Fig. 5G**). These data collectively show that both RmcA and MorA physically interact with PelD, but do so maximally under conditions where other components of the Pel synthase complex are present alongside PelD (39), as illustrated in **Figure 5H**.

## Discussion

Exploiting previous findings from our lab in which a CRISPR-activated strain exhibited a defect in biofilm maintenance (21), we discovered that two PDEs, RmcA and MorA, were essential for maintaining late-stage biofilms. The Δ*rmcA* and Δ*morA* mutants exhibit phenotypes consistent with the inability to degrade c-di-GMP, specifically, elevated c-di-GMP, increased Pel production and the ability to initiate a robust biofilm. Yet, the Δ*rmcA* and Δ*morA* mutants fail to maintain the biofilm in long term static assays or when established biofilms are deprived of a carbon source in a microfluidic chamber. Additionally, we have shown that the inability to maintain biofilms in these mutant backgrounds is driven by widespread cell death during nutrient limitation. Consistent with the hypothesis that cell death in these mutants is due to an aberrant nutrient limitation response, we showed that the Δ*relA* Δ*spoT* mutant, which lacks the ability to induce a stringent response, demonstrates a biofilm maintenance defect during nutrient limitation similar to that observed for the Δ*rmcA* and Δ*morA* mutants.

Taken together, the above data suggests a model (**Fig. 6**) whereby the production of the energetically expensive Pel polysaccharide, required for the initial steps of biofilm formation, is downregulated by RmcA and MorA during biofilm maintenance when nutrient limitation conditions predominate. As such, while the loss of either PDE results in increased EPS and enhanced biofilm growth, a boon to these microorganisms in resource-rich environments typical of early biofilm formation, it leaves the cells unable to adapt to later nutrient-limited conditions in the context of a mature biofilm.

**Figure 6.**
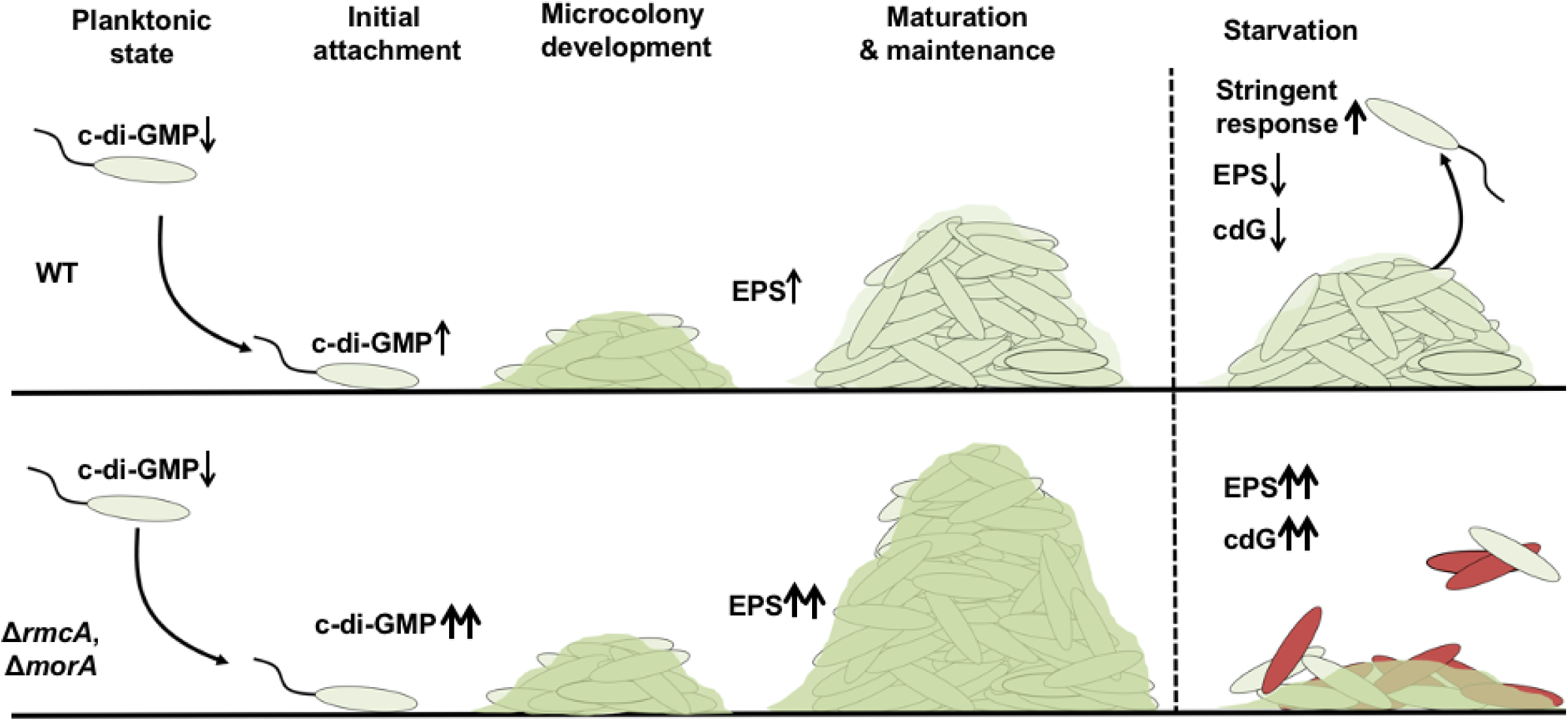
A model for MorA and RmcA-mediated biofilm maintenance. Model for RmcA- and MorA-mediated biofilm maintenance in *P. aeruginosa*. Typical biofilm development (top panel) involves surface attachment, after which increased c-di-GMP and EPS synthesis mediate microcolony development and increased biofilm biomass. During the maturation and maintenance phase of biofilm development, regulatory changes reflect growth in a nutrient-limited environment and result in a decrease in the production of energetically expensive products like Pel, reduced c-di-GMP (cdG) and induction of the stringent response. Loss of RmcA or MorA (bottom panel) results in enhanced c-di-GMP and Pel production and increased biomass in nutrient rich environments. The Δ*rmcA* and Δ*morA* mutant biofilms are unable to appropriately respond to nutrient limitation, resulting in cell death and loss of biofilm biomass.

The mechanisms by which RmcA and MorA are regulated in nutrient-limited conditions remain unknown, however recent findings within *Pseudomonas putida* provide a potential signaling framework. Work by Carlos Díaz-Salazar et al. found that RelA and SpoT-dependent synthesis of (p)ppGpp mediates dispersal during nutrient-limited conditions (40). This group also found that (p)ppGpp increased transcription of the PDE *bifA* and that a Δ*bifA* mutant was defective in starvation-induced biofilm dispersal (40, 41). It is possible that RmcA and MorA within *P. aeruginosa* operate analogously to that of BifA in *P. putida* by acting as effectors of stringent response signaling during nutrient-limited conditions. Unlike BifA in *P. putida*, RmcA and MorA do not coordinate dispersal in *P. aeruginosa*, but rather participate in effectively maintaining the biofilm in the face of nutrient limitation.

The high levels of c-di-GMP in the Δ*rmcA* and Δ*morA* mutants may have adverse impacts on the cells in nutritionally-limited, mature biofilms. While we do not yet completely understand how the regulation of Pel may contribute to biofilm maintenance, there is strong evidence for the physical interaction of RmcA and MorA with PelD and other elements of the Pel biosynthetic machinery (**Fig. 5E-H**), suggesting that RmcA and MorA may have a direct role in regulating Pel synthesis. However, as noted above, artificially increasing Pel expression to high levels does not result in increased cell death in nutrient-limited condition (**Fig. S7B**), suggesting that the overexpression of Pel alone is insufficient to drive the late-stage cell death phenotype of the Δ*rmcA* and Δ*morA* mutants.

Further studies are needed to elucidate whether the potential role of Pel in biofilm maintenance is related to the stringent response. It is possible that the inability to appropriately regulate c-di-GMP and EPS production during nutrient limitation impacts (p)ppGpp levels, eventually resulting in extensive cell death and biofilm degradation. Alternatively, the ability to induce a stringent response may be part of a coordinated down-regulation of metabolic activity required for the long-term maintenance of a mature biofilm, particularly when carbon/energy sources are limiting. Additionally, Pseudomonads appear to have developed catabolic pathways for the utilization of arginine and lactate for “maintenance energy” in mature biofilms (43, 44), as well as a pathway to down-regulate flagellar motility, another early-stage biofilm factor, in mature biofilms (1). Taken together, these data indicate that Pseudomonads, and likely other microbes, have active, well-regulated mechanisms necessary to maintain a mature biofilm in the face of changing environmental conditions.

Finally, the apparent role for c-di-GMP-metabolizing enzymes RmcA and MorA later in the biofilm lifestyle suggests the interesting possibility that the plethora of these enzymes in Pseudomonads stems from their roles in regulating discrete aspects of the biofilm lifestyle – from formation to maturation to maintenance to dispersal. The finding that the loss of different PDEs would result in different phenotypes may be expected given the varied impacts that these enzymes have on the regulation and timing of c-di-GMP signaling and biofilm formation (9, 13). Here, while we assessed all PDE and dual-domain mutants in *P. aeruginosa* in our initial screen, we only observed consistent and significant defects in biofilm maintenance in the Δ*rmcA* and Δ*morA* mutants. Previous work identified a number of DGCs and PDEs apparently required for early biofilm formation, including SadC, RoeA, BifA, and SiaD (15, 16, 45). In contrast, the PDE DipA has been shown to mediate biofilm dispersion in response to elevated nutrient concentrations and this protein localizes to the cell pole during division, resulting in the asymmetric distribution of c-di-GMP (18, 46). Thus, our data are consistent with the hypothesis of stage-specific roles for DGCs/PDEs in the biofilm life cycle.

The network which controls c-di-GMP levels in *P. aeruginosa* is complex. Identified first in *P. putida*, MorA was found to repress motility in swim assays (37). This same work found that the enhanced motility of the *morA* mutant was not observed in *P. aeruginosa* (37), but previous work from the Hogan and O’Toole labs showed that the Δ*morA* mutant exhibited a significant decrease in flagella-dependent swimming and swarming motility (24). The basis of this difference in phenotypes may be due to the fact that different species of *Pseudomonas* were used in the two studies. Nevertheless, given the role of motility in early biofilm formation, the observation that MorA contributes to swimming and swarming motility indicates that this PDE also likely contributes to the initiation of biofilm communities.

Insightful work from the Dietrich lab provided evidence that RmcA is activated by phenazine availability to mediate a decrease in c-di-GMP levels during oxidative stress conditions (25). In this model, RmcA can act as a redox sensor and may behave as a switch to translate this signal into decreased levels of c-di-GMP and EPS. This model for the role of RmcA in the context of the colony biofilm used by Okegbe, Dietrich and colleagues (25) is largely in agreement with the experimental evidence we have provided, which suggests that RmcA is important for biofilm maintenance. Specifically, it is likely that in mature biofilms with elevated biomass, nutrient limitation coincides with oxygen depletion and the production and utilization of phenazines as electron shuttles. Indeed, the direct regulatory signal sensed by RmcA appears to be a change in redox state that is likely secondary to the loss of a catabolizable carbon source (25). Thus, in this environment, we hypothesize that RmcA downregulates the production of energetically-expensive Pel EPS, and that failure to do so could result in the observed cell death and biofilm maintenance defect. Together, these data suggest that further examination of these enzymes will generate a more nuanced view of the model presented in **Figure 6**, wherein specific c-di-GMP metabolizing enzymes work at one or more stages of the biofilm life cycle, with the potential to perform several overlapping functions across these various stages (i.e., biofilm initiation and biofilm maintenance).

## Materials and Methods

### Strains and media

Strains, plasmids and primers used in this study are listed in supplemental Tables S1 and S2. *P. aeruginosa* strain UCBPP-PA14 (PA14) was used in all the experiments, except for the IP studies and the proteins used in the BATCH, which used the PAO1 strains. *P. aeruginosa* were routinely streaked onto lysogeny broth (LB) plates containing 1.5% agar prior to overnight culturing in LB liquid cultures at 37°C. When appropriate, LB was supplemented with 10 µg/ml gentamicin (Gm) and 50 µg/ml kanamycin (Kan). Biofilm medium used in static 96 well crystal violet assays was composed of M63 medium supplemented with 1 mM MgSO_4_ and 0.4% (w/v) L-arginine monochloride, and where indicated the arginine was replaced by 20 mM pyruvate (3, 47). ALI and microfluidic-based biofilm assays were performed in KA medium, a modification of the previously reported K10T medium (48) containing 50 mM Tris-HCl (pH 7.4), 0.61 mM MgSO_4_ and 0.4% arginine.

### Static biofilm assays and quantification

Overnight cultures were inoculated into 96-well U-bottom polystyrene plates (Costar) containing M63-based medium and grown for the specified time at 37°C in a hydrated container prior to washing, staining with crystal violet, and then solubilization of the crystal violet stain with 30% glacial acetic acid (3, 47). Biofilm was quantified by measuring the extent of biofilm-associated CV solubilized in a spectrophotometer at OD_550_.

For biofilms imaged microscopically along the ALI, overnight cultures were prepared as described above and inoculated into 12-well dishes containing KA biofilm medium. Glass coverslips were partially submerged in the medium and grown at 37°C for the desired length of time prior to removal and imaging with a Nikon Eclipse Ti inverted microscope where a minimum of ten fields of view were captured. To assess viability of coverslip-grown biofilms, propidium iodide and Syto9 (Molecular Probes® Live/Dead BacLightTM) were gently mixed into the growth medium and biofilms were stained for one hour prior to imaging. Pel was visualized within the ALI with the addition of 10 µl/ml of Pel-specific fluorescein-labelled *Wisteria floribunda* lectin (Vector Laboratories) to the KA medium at the start of the assay. To assess the concentration of c-di-GMP we utilized the P_*cdrA*_-*gfp* fusion expressed from a multicopy pMQ72 (49) which was maintained in overnight cultures supplemented with Gm prior to inoculation into antibiotic-free KA.

### Congo Red assay

Strains were grown overnight in 5 ml LB at 37°C and 5 µl of an overnight culture was spotted onto plates containing 1.5% agar, 1% Tryptone, 40 μg/mL Congo Red (CR) and 15 μg/mL Coomassie brilliant blue. The plates were incubated at 37°C for 24 h and imaged after an additional 4 days at room temperature.

### Microfluidics

Biofilms were visualized under flow in microfluidics chambers kindly provided by the Nadell laboratory. Chambers used poly-dimethylsiloxane (PDMS) bonded to a 1.5 × 36mm X 60mm cover glass (ThermoFisher, Waltham MA) through soft lithography techniques (50, 51). Overnight bacterial cultures were centrifuged, resuspended in KA, adjusted to an OD_600_ of 1, pipetted into microfluidics chambers and allowed to attach for 1h. Tubing (#30 Cole Palmer PTFE) to transport influent and effluent medium was attached first to BD 5-ml syringes containing KA biofilm medium, then to the microfluidics chambers and then to syringe pumps (Pico Plus Elite, Harvard Apparatus) operating at a flow rate of 0.5 µl/min.

### Image acquisition and data analysis

All microscopy was acquired using Nikon Elements AR running a Nikon Eclipse Ti inverted microscope equipped with a Hamamatsu ORCA-Flash 4.0 camera and imaged through either a Plan Apochromat 100x DM Oil or Plan Fluor 40x DIC M N2 objective. Fast scan mode and 2×2 binning was used for imaging. All images were collected in a temperature controlled environmental chamber set to 37°C. Images were processed with background subtraction and signal strength quantified by measuring mean signal intensity/pixel through the Integrated Density (IntDen) function.

### Statistical analysis

Data was analyzed with Graph Pad Prism 8. Unless otherwise noted, data are representative of the results from at least three independent experiments. A Student’s *t* test was used to compare results and to assess significance.

### Strain construction of fluorescent strains

*P. aeruginosa* expressing fluorescent GFP were made through electroporation of a multi-copy plasmid pSMC21. mKO was introduced in single copy on the chromosome at the *att*:Tn*7* site via conjugation from *E. coli* S17-λpir pCN768 (52). In-frame, unmarked *rmcA* and *morA* gene deletions were generated using allelic replacement, as reported (53). Construction of gene deletion alleles was performed by amplifying flanking regions of the *rmcA* and *morA* ORFs and joining these flanking regions by splicing-by-overlap extension PCR. The upstream forward and downstream reverse primers were tailed with restriction endonuclease cleavage sequences to enable ligation-dependent cloning of the spliced PCR products. The assembled Δ*rmcA* and Δ*morA* alleles were ligated into pEX18Gm (54) and the resultant allelic exchange vectors were transformed into *E. coli* DH5α. Plasmids were then isolated from individual colonies and verified by Sanger sequencing using M13F and M13R. Plasmids were conjugated into *P. aeruginosa* PA14 from E. coli and merodiploids selected on LB agar containing 10 μg/mL gentamicin. SacB-mediated counter selection was carried out to select for double crossover mutations on no salt LB (NSLB) agar containing 15% (w/v) sucrose. Unmarked gene deletions were identified by colony PCR using primers that targeted the outside, flanking regions of the *rmcA* and *morA* ORFs. These PCR products were Sanger sequenced using the same primers to confirm the correct deletion.

The Δ*rmcA* and Δ*morA* deletion alleles were introduced into *P. aeruginosa* PAO1 Δ*wspF* Δ*psl* P_BAD_*pel* (38) via biparental mating with the donor strain *E. coli* SM10 (55). Merodiploids were selected on Vogel-Bonner minimal medium (VBMM) agar containing 30 μg/mL gentamicin. SacB-mediated counter selection was performed to select for double crossover mutations on no salt LB (NSLB) agar containing 15% (w/v) sucrose. Unmarked gene deletions were identified by colony PCR using primers that targeted the outside, flanking regions of the *rmcA* and *morA* ORFs. These PCR products were Sanger sequenced using the same primers to confirm the correct deletion.

For gene complementation in *P. aeruginosa*, pUC18T-miniTn7T-Gm, which allows for single-copy chromosomal insertion of genes (56), was modified to allow for arabinose-dependent expression of complementing genes. The *araC*-P_BAD_ promoter from pJJH187 (57) was amplified using the primer pair miniTn7-pBAD-F and miniTn7-pBAD-R, the latter of which contains flanking sequence encoding *Sma*I, *Not*I, *Pst*I, and *Nco*I sites to generate a multiple cloning site downstream of the *araC*-P_BAD_ promoter. The resulting PCR product was cloned into the *Sac*I and *Hin*dIII sites of pUC18T-miniTn7T-Gm to generate pUC18T-miniTn7T-Gm-pBAD.

The ORF corresponding to *rmcA* or *morA* was amplified using primer pairs tailed with restriction endonuclease cleavage sequences to enable ligation-dependent cloning of the PCR products. Upstream primers were also tailed with a synthetic ribosome binding site upstream of the start codon. PCR products were ligated into pUC18T-miniTn7T-Gm-pBAD and the resulting miniTn*7* vectors were transformed into *E. coli* DH5α and selected on LB agar containing 10 μg/mL gentamicin and 100 μg/mL carbenicillin. Plasmids were then isolated from individual colonies and verified by Sanger sequencing using the miniTn7-SEQ-F and miniTn7-SEQ-R primers, as well as primers specific to internal portions of each gene, as appropriate.

Incorporation of C-terminal vesicular stomatitis virus glycoprotein (VSV-G) tags into p-miniTn7-*rmcA* and p-miniTn7-*morA* was performed via PCR with 5’-phosphorylated primer pairs. The forward primer annealed to the stop codon of *rmcA* or *morA* plus 15-22 bp of downstream vector encoded sequence. The reverse primer annealed to the coding strand 19-21 bp upstream of the *rmcA* or *morA* stop codon. The forward and reverse primers contained 5’-overhangs that encoded the last and first halves, respectively, of the VSV-G peptide sequence. The PCR amplified product of these primer pairs was subsequently digested with *Dpn*I for 1 h at 37 °C to remove template DNA, followed by incubation with T4 DNA ligase overnight at room temperature to self-ligate the blunt ends and re-circularize the vector. The resulting C-terminally VSV-G-tagged miniTn7 vectors were transformed into *E. coli* DH5α and selected on LB agar containing 10 μg/mL gentamicin and 100 μg/mL carbenicillin. Plasmids were then isolated from individual colonies and verified by Sanger sequencing as described above.

Complemented *P. aeruginosa* strains were generated through incorporation of miniTn7 vectors at the neutral *att::*Tn*7* site on the *P. aeruginosa* chromosome via electroporation of miniTn7 vectors, along with the helper plasmid pTNS2, as previously described (56). Transposon mutants were selected on LB agar containing 30 μg/mL gentamicin.

### Co-immunoprecipitation assays

1 L of LB, containing 0.5% (w/v) L-arabinose and 30 μg/mL gentamicin, was inoculated with a *P. aeruginosa* strain carrying a VSV-G-tagged protein and allowed to grow overnight at 37 °C with shaking at 200 RPM. The next day, cells were collected at 5,000 × *g* for 20 min at 4 °C. Cell pellets were resuspended in 5 mL of 0.2 M Tris-HCl pH 8, 1 M sucrose, 1 mM EDTA, and 1 mg/mL lysozyme. Cells were incubated for 10 min at room temperature prior to the addition of 20 mL of water and further incubation on ice for 30 min. The resultant solution was centrifuged at 30,000 × *g* for 20 min at 4 °C to collect spheroplasts. The pellet was then resuspended in 50 mL of 10 mM Tris-HCl pH 7.5, 5 mM EDTA, and 1 mM DTT, and lysed by homogenization using an Emulsiflex-C3 (Avestin Inc.) at a pressure of 10,000 - 15,000 psi until the solution appeared translucent. The solution was clarified by centrifugation at 30,000 × *g* for 20 min at 4 °C, and the resultant supernatant was further centrifuged at 180,000 × *g* for 1 h at 4 °C to collect the membranes. Membranes were then solubilized in 10 mL of Buffer A (50 mM Tris-HCl pH 8, 10 mM MgCl_2_, and 2% (w/v) Triton X-100) using a Dounce homogenizer and centrifuged at 90,000 × *g* for 30 min at 4 °C to clarify the solution. A sample of the solubilized membranes was collected before application to the IP resin as a representative example of the input into the experiment. The IP resin (Sigma anti-VSV-G monoclonal antibody-agarose conjugate) was prepared by mixing 60 μL of resin slurry with 10 mL of Buffer A, followed by collection of the IP resin by centrifugation at 100 × *g* for 2 min at 4 °C and removal of the supernatant. The solubilized membranes were applied to the washed IP resin and incubated at 4 °C for 1 h with agitation. The IP resin was then collected by centrifugation at 100 × *g* for 2 min at 4 °C and the supernatant discarded. The resin was washed five times with 10 mL of Buffer A as above to remove non-specifically bound protein. The resin was then mixed with 150 μL of 2× Laemmli buffer, boiled at 95 °C for 10 min, and analyzed by SDS-PAGE followed by Western blotting as described below. As a negative control, the above experimental protocol was also performed using a *P. aeruginosa* strain carrying the corresponding untagged protein.

### Western blot sample analysis

For Western blots, a 0.2 μm PVDF membrane was wetted in methanol and soaked for 5 min in Western transfer buffer (25 mM Tris-HCl, 150 mM glycine, 20% (v/v) methanol) along with the SDS-PAGE gel to be analyzed. Protein was transferred from the SDS-PAGE gel to the PVDF membrane by wet blotting (25 mV, 2 h). The membrane was briefly washed in Tris-buffered saline (10 mM Tris-HCl pH 7.5, 150 mM NaCl) containing 0.5% (v/v) Tween-20 (TBS-T) before blocking in 5% (w/v) skim milk powder in TBS-T for 2 h at room temperature with gentle agitation. The membrane was briefly washed again in TBS-T before incubation overnight with primary antibody (1:5000 α-PelD polyclonal antibody (36) or 1:75000 Sigma α-VSV-G monoclonal antibody) in TBS-T with 1% (w/v) skim milk powder at 4 °C. The next day, the membrane was washed four times in TBS-T for 5 min each before incubation for 1 h with secondary antibody (1:2000 dilution of BioRad affinity purified goat α-rabbit or goat α-mouse IgG conjugated to alkaline phosphatase) in TBS-T with 1% (w/v) skim milk powder. The membrane was then washed four times with TBS-T for 5 min each before development with 5-bromo-4-chloro-3-indolyl phosphate/nitro blue tetrazolium chloride (BioShop ready-to-use BCIP/NBT solution). Developed blots were imaged using a BioRad ChemiDoc imaging system.

### Bacterial adenylate cyclase two-hybrid (BACTH) assays

Cloning of *rmcA* and *morA* into the BACTH assay-compatible vector pUT18C, and *pelD* into pKT25, was performed using standard molecular methods. Reverse primers were flanked with a 3’-stop codon for cloning into pUT18C and pKT25, which encode a N-terminal adenylate cyclase fragment fusion (39). Primer pairs were tailed with restriction endonuclease cleavage sequences to enable ligation-dependent cloning and were used to amplify the corresponding *pelD, rmcA*, and *morA* genes from PAO1 genomic DNA. PCR products were digested with the appropriate restriction endonucleases and ligated into pUT18C and pKT25, as appropriate. Ligations were transformed into *E. coli* DH5α and selected on LB agar containing 50 μg/mL kanamycin for pKT25 clones, or 100 μg/mL carbenicillin for pUT18C clones. Plasmids were then isolated from individual colonies and verified by Sanger sequencing using primers specific for pUT18C and pKT25, as well as primers specific to internal segments of *rmcA* and *morA*, as appropriate. Positive clones were verified as above.

Combinations of the above T18 and T25 fusion proteins were transformed into the BACTH compatible strain BTH101 (Euromedex) for analysis. For each experiment, 5 mL of LB supplemented with 50 μg/mL kanamycin, 100 μg/mL carbenicillin, and 0.5 mM IPTG was inoculated with the appropriate BTH101 strain and grown overnight at 30 °C with shaking at 200 RPM. The next day, 2 μL of culture was used to spot inoculate a LB agar plate containing 50 μg/mL kanamycin, 100 μg/mL carbenicillin, 0.5 mM IPTG, and 50 μg/mL 5-bromo-4-chloro-3-indolyl-β-D-galactopyranoside (X-Gal). The plates were incubated for 24 h at 30 °C and subsequently photographed. The vectors pUT18C::zip and pKT25::zip (39) were used as a positive control. Empty pUT18C and pKT25 vectors were used as a negative control.

To generate tag-free expression constructs for the modified BACTH assays, *pelE* and *pelG* were amplified from PAO1 genomic DNA using forward primers that were flanked with a synthetic ribosome binding site and reverse primers flanked with a 3’-stop codon. PCR products were subsequently digested with *Eco*RI and *Hin*dIII and ligated into the arabinose-inducible expression vector pBADGr (10 Clones with positive inserts were verified by Sanger sequencing using the BADGr-SEQ-F and BADGr-SEQ-R primers). The vector expressing both *pelE* and *pelG* was generated by amplifying the *pelE* and *pelG* ORFs from *P. aeruginosa* genomic DNA. The intervening *pelF* ORF from the *pel* operon was excluded by joining the upstream *pelE* and downstream *pelG* ORFs via the splicing-by-overlap extension PCR method, as described above for generation of chromosomal deletion alleles, to generate a single polycistronic strand. All assays with untagged constructs were performed as above, with the addition of 10 μg/mL gentamicin and 0.5% (w/v) arabinose to all growth media for, respectively, maintenance of and expression from pBADGr. Empty pBADGr was used as a vector control.

## Acknowledgements

We thank C. Nadell for providing the fluorescent protein constructs and help with preparing the microfluidic chambers. We thank A. Collins and G. Heussler for the constructive feedback throughout the course of this research. This work was supported by NIH Grant R37 AI83256-06 to G.A.O and the Canadian Institutes of Health Research (CIHR) MOP 43998 and FDN154327 to PLH. PLH is the recipient of a Tier I Canada Research Chair. GBW was supported in part by a Canada Graduate Scholarship from the Natural Sciences and Engineering Research Council of Canada (NSERC) and a graduate scholarship from Cystic Fibrosis Canada.

## List of Supplementary Material

Figure S1. The biofilm maintenance defect in PDE mutants is independent of carbon source.

Figure S2. Elevated c-di-GMP does not in increase the biofilm deficit in the Δ*morA* or Δ*rmcA* mutants.

Figure S3. The biofilm maintenance defect can be partially rescued in the static assay with fresh medium.

Figure S4. Peroxide addition does not induce biofilm defect.

Figure S5. Cell death during late-stage biofilm differs between *ΔrmcA* and *ΔmorA*.

Figure S6. Initial biofilm formation by the WT and Δ*rmcA* and Δ*morA* mutants.

Figure S7. Viable count of the WT and PDE mutants in the microfluidic device effluent.

Figure S8. Late stage biofilm defect cannot be induced or rescued with changes to Pel concentration.

Table S1. Bacterial strains and plasmids used in this study.

Table S2. List of Primers.

